# 3D protein structure from genetic epistasis experiments

**DOI:** 10.1101/320721

**Authors:** Nathan J. Rollins, Kelly P. Brock, Frank J. Poelwijk, Michael A. Stiffler, Nicholas P. Gauthier, Chris Sander, Debora S. Marks

## Abstract

High-throughput experimental techniques have made possible the systematic sampling of the single mutation landscape for many proteins, defined as the change in protein fitness as the result of point mutation sequence changes. In a more limited number of cases, and for small proteins only, we also have nearly full coverage of all possible double mutants. By comparing the phenotypic effect of two simultaneous mutations with that of the individual amino acid changes, we can evaluate epistatic effects that reflect non-additive cooperative processes. The observation that epistatic residue pairs often are in contact in the 3D structure led to the hypothesis that a systematic epistatic screen contains sufficient information to identify the 3D fold of a protein. To test this hypothesis, we examined experimental double mutants for evidence of epistasis and identified residue contacts at 86% accuracy, including secondary structure elements and evidence for an alternative all-α-helical conformation. Positively epistatic contacts – corresponding to compensatory mutations, restoring fitness – were the most informative. Folded models generated from top-ranked epistatic pairs, when compared with the known structure, were accurate within 2.4 Å over 53 residues, indicating the possibility that 3D protein folds can be determined experimentally with good accuracy from functional assays of mutant libraries, at least for small proteins. These results suggest a new experimental approach for determining protein structure.

## Introduction

Experimental work dating back over 20 years has shown that epistasis, non-additive phenotypes of combined mutations, within a protein sometimes occur between residues that are spatially close in that protein’s 3D structure(s). The reported correspondence between physical proximity and non-additive effect was mostly for compensatory epistasis - where the effect of two mutations is less deleterious than expected from the combined effect of the single mutations (*1–3*). Until recently, studies of epistasis have been limited to measuring relatively few mutant pairs *(1–3)*, but recent investigations have taken advantage of technological advances in sequencing ability to assess the phenotypic effects of thousands of mutations simultaneously, generally coined “deep mutational scanning” *(4–29)*. These experiments can measure different phenotypes including ligand binding, splicing and catalysis *(5, 9, 12, 14, 22, 24, 30)*, and cellular or organismal fitness under selection pressure *(6–8, 10, 13, 15, 18, 20, 21)*. Typically, these scans survey all possible single mutations, with a few that have surveyed a large number (but small minority) of multiple mutations *(7, 23)*, and one study assessed the effects of all double mutations across an entire protein domain: the 56-residue IgG binding domain of protein G from Streptococcus (GB1) (*31*). Confirming many earlier lower throughput studies, Olson et al. found that epistatic pairs of residues in the GB1 domain of protein G have a tendency to be close in 3D, especially those that have positive, compensatory epistasis (*31*).

Meanwhile, structural biology has also benefited from advances in sequencing. In addition to experimental determination (e.g. NMR, X-Ray crystallography and cryo-EM), recent computational approaches have also been successful in folding both small and large proteins *ab initio* using residue-residue contacts inferred from natural, evolutionary sequence information (*32–36*). These successful methods use a global model, e.g., inferred from entropy maximization to identify residue-residue evolutionary couplings also likely to be close in the protein’s folded structure. Where sufficient related sequences are available, evolutionary couplings provide enough information to determine accurate 3D structures for many protein examples, including those of large membrane proteins (*32, 33*), complexes (*37*), and RNAs (*38*), as well as proteins with alternative conformations (*39*). These new computational methods, although powerful, are limited by the availability of large and diverse sequence families from the natural environment.

By analogy to the success of computational approaches determined from natural sequence variation, we reasoned that the strongest epistatic pairs of residues determined from large-scale genetic experiments may be sufficient to fold a protein into its native, functional three-dimensional structure. Here we test this idea by computing the epistasis of all possible double mutants as reported by Olson et al. (*31*), who assayed more than 500,000 sequences for fitness in a binding assay. We find that the positional pairs that contain the strongest compensatory epistatic effects in GB1 are sufficient to reveal its secondary structural elements and its native three-dimensional fold, with local epistatic contacts also signaling a possible alternative secondary structure consistent with the triple-helical bundle as demonstrated in prior fold-switching studies *(40–43)*. These results indicate that deep mutational scans can provide a new method of assaying protein structure.

## Results

### Residue pairs that are epistatic are close in 3D

To explore whether pairs of residues that are most epistatic are close in the 3D structure, we analyzed the experimental data from Olson et al. *(31)* that systematically tested all single and almost all pairwise mutations of the 56 amino acid GB1 domain of Streptococcal protein G (1045 singles and 535,917 double mutations). In these experiments, the pooled proteins were assayed for binding to human immunoglobulin IgG, and fitness was approximated by changes in sequence read ratios using mRNA tags on the proteins. We filtered out reads with the highest probability of noise, discarding 3% of the double mutants (those with fewer than 20 reads before selection). We then estimated the epistasis for each double mutant using a correction for the assay resolution, reproducing the epistasis scores computed in the original report *(31)* (Supp. Figure 1, Supp. Table 1, Methods). We then mapped the pairs of epistatic residues onto known three-dimensional structures of the GB1 protein, i.e., the 14 solved 3D structures of wild-type GB1 (Figure 1, Supp. Table 2). 95% of the top 20 (86% of the top L/2 pairs where L = sequence length) of residues contain any atoms within 5Å of each other in the NMR structure 2gb1(44) (Table 1), with similar precisions when compared against other wildtype crystal structures (Supp. Table 2) *(45)*. We observed compensatory epistasis (positive epistasis) to be more informative than negative epistasis for 3D structure contacts (*31*) by comparing positive and positive epistatic pairs to contacts in known structures (Supp. Table 2, Methods). Here, we report results for contacts predicted by the largest positive, negative, and absolute epistasis for comparison.

**Figure 1.**
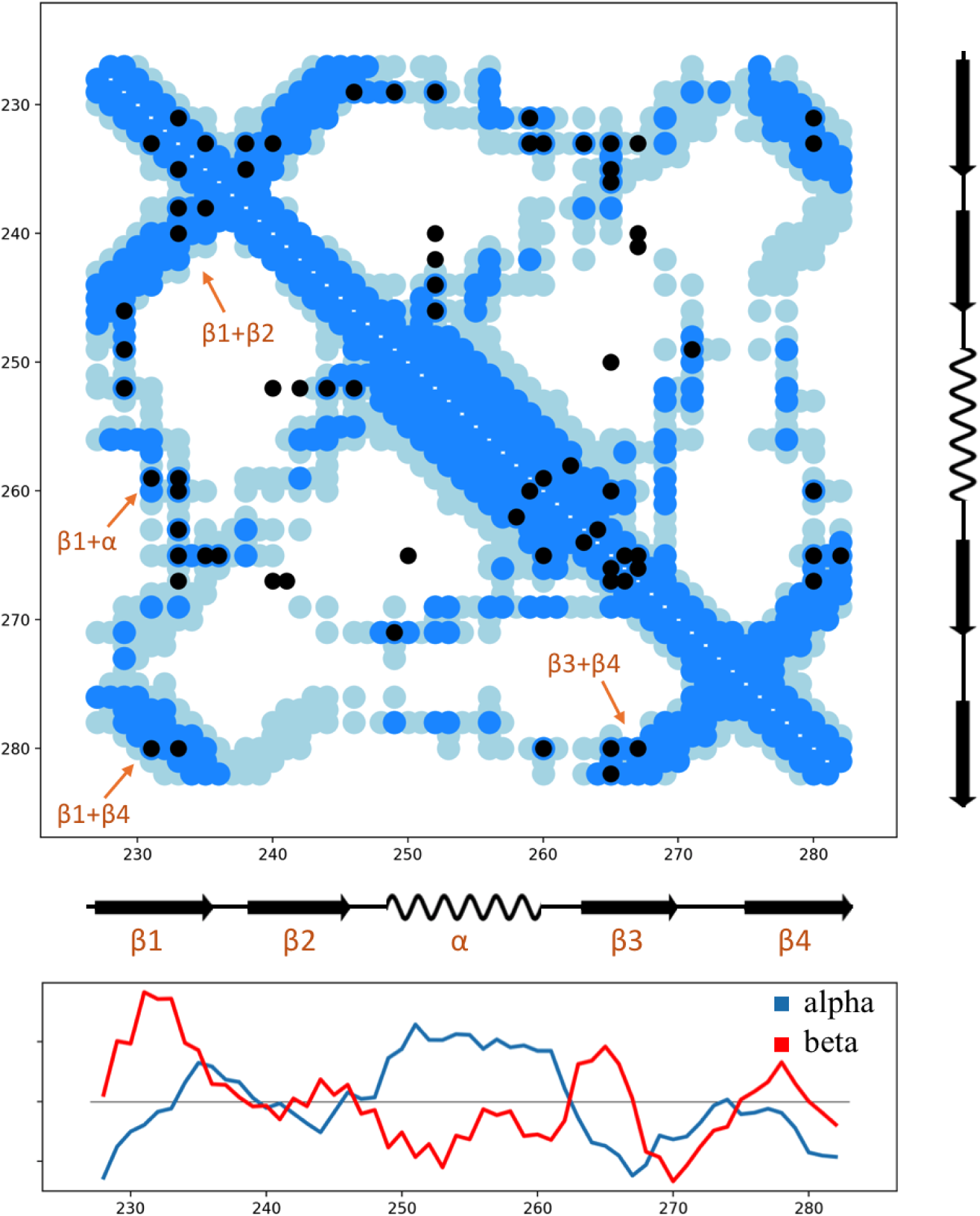
Amino acid residue contact prediction of the GB1 protein domain from epistatic couplings. **Top:** Top positive epistatic pairs (black) up to L/2 long-range (|i-j| ≥ 5) contacts (L: length of protein, 37 contacts total, 83% true positives) that are used to determine the topological arrangement of secondary structure elements (orange arrows). Contact predictions are precise where the epistatic pair (black) agrees with the contacts in the crystallographic structure (minimum atom-atom distance between two residues; dark blue, 5 Å cutoff; light blue, 8 Å cutoff) and less precise or incorrect otherwise. **Bottom:** Secondary structure propensity from local epistasis values (vertical axis): α helical score (blue), β strand score (red). Scores are mean centered with arbitrary units (values in Supp. Table 3).

**Table 1.** Percentage of correctly predicted contacts (true positives) using various forms of epistasis. The percent of predicted contacts according to residues with any atom within 5 Å (NMR structure 2gb1). Three forms of epistasis are considered for prediction: i,j pairs corresponding to mutations with the largest positive, negative, or absolute measure epistasis.

|  | *top 20 pairs* | *Top 30 pairs* | *top 40 pairs* | *top 50 pairs* | *top 100 pairs* |
| --- | --- | --- | --- | --- | --- |
| *Positive epistasis* | 95% | 87% | 83% | 72% | 62% |
| *Negative epistasis* | 25% | 27% | 30% | 38% | 31% |
| *Absolute epistasis* | 40% | 40% | 40% | 44% | 37% |

### Epistasis scores between nearby residues reveal secondary structure

By analogy to previous work using evolutionary sequence alignments to determine secondary structure *(39)*, we used the strength of the epistasis between residues i to i+1, i+2, i+3, and i+4 to explore the secondary structure of the domain. This approach using only information contained has the advantage of being unsupervised and capable of revealing alternative states (in contrast to the successful existing methods available by training from known structures, e.g., PredictProtein *(46)* and PsiPred (*47*)). The secondary structure we computed from the local epistasis scores is almost identical to those in known structures of GB1, a four-stranded β-sheet with one α-helix, differing only in an offset at the start of the second and third β-strands and a weaker signal for the second β strand compared to the other three (Figure 1). Visualization of the interactions of these elements also reveals typical secondary structure element interactions, such as long-range (in sequence) strand-strand contacts (Figure 1). From the contact maps of the most positive epistatic pairs, we observe two antiparallel β-strand hairpin contacts (strands β1-β2 and strands β3-β4) and one parallel β-strand contact (strands β1-β4), consistent with a β sheet with strand order β2-β1-β4-β3 (Figure 1). These predicted epistatic pairs define the strand-strand register to within +/-1 residue and provide insight into the overall 3D topology of the protein.

Our analysis also reveals a signal for alternative secondary structural elements, with a propensity towards two helices in positions 235-240 and 268-280 in addition to the existing middle helix in the GB1 domain. This competing signal suggests three consecutive helices as an alternative to the four β-strands plus one helix secondary structure of GB1 (Figure 1, Supp. Table 3). Intriguingly, the three helices match the secondary structure of a mildly mutated GB1 structure with different overall 3D fold *(40–42)*, which is topologically similar to the structure of the GA domain of the same protein that binds human serum albumin - a different partner *(43)*. However, this signal seems specific to local epistasis. By contrast, there are a small number of high-scoring evolutionary couplings (derived from natural sequences) that are uniquely consistent with the tertiary structure of the three α-helical fold. Hence it seems that the evolutionary sequence evidence more strongly reveals a potential three α-helical fold than the double mutation experiments. This could be explained by the nature of the experimental selection to bind IgG rather than serum albumin. It seems that more research is needed both to tease out how the fold switching might be encoded in the sequence and to understand its biological relevance.

### Three-dimensional folds from epistasis experiments alone

We then tested the efficacy with which epistasis-predicted contacts can guide folding towards correct three-dimensional structure. Using 10 to sequence length 56 of the most positively epistatic pairs as residue-residue distance constraints, as well as secondary structure constraints, we folded the open chain according to a distance geometry and simulated annealing protocol in the “Crystallography and NMR software Suite”, CNS *(48)* (Methods). We used additional rounds of simulated annealing in CNS to further refine structures. Resultant 3D models were ranked by the number of satisfied residue-residue constraints and realistic distances between beta-strands (Supp. Figure 2, Methods). The top-ranked model is 2.4 Å C-α rmsd over 53 residues (TM-align score of 0.57) compared to 2gb1 (*45, 49*), and therefore in excellent overall agreement with the known experimental structure (Figure 2). Removing 4 false positives from the 28 (L/2) contacts further improves the folding to 1.9 Å C-α rmsd, which unsurprisingly suggests that folding can be improved by better detecting epistatic pairs not likely to be in direct contact. These false positives may contain information relevant to biological function; all four pairs involve residues at the immunoglobin-binding interface (Figure 2). Additionally, these models are likely to be improved by adding additional strand-strand constraints (as in (*33*)).

**Figure 2.**
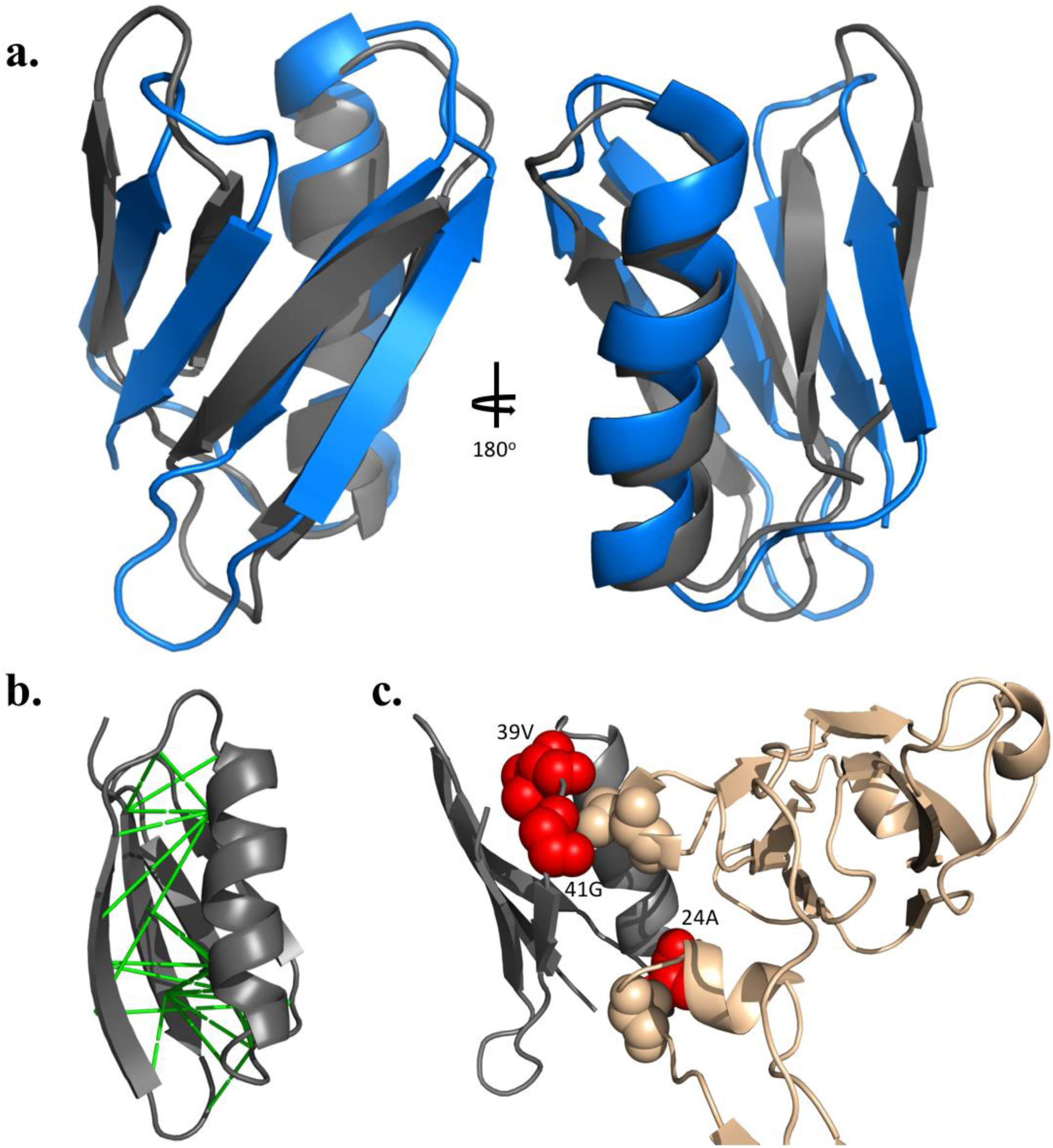
Predicted 3D fold model of the GB1 protein domain. (gray) generated from positive epistatic couplings, compared to the crystal structure (**a**, PDBid: 2gb1 in blue). The predicted structure is within 2.4 Å of the known structure over 53/56 residues, with a template modeling (TM) score of 0.57. Top-scoring epistatic pairs (**b**, green lines on 2gb1) are used as constraints to fold the protein. However, four of these epistatic pairs are not close within the known structure, and interestingly all involve sidechains that contact immunoglobin when bound (**c**, GB1 in grey with red highlighted residues, IgG in tan, PDBid: 1fcc (*54*)).

### Comparison to evolutionary couplings from natural evolution

We then examined whether evolutionary couplings, derived from the alignment of natural sequences alone, provide overlapping or orthogonal information to epistatic-derived contacts from *in vitro* fitness assay experiments. Obtaining high-quality evolutionary couplings relies on a protein having a traceable evolutionary history large enough to provide many related sequences that are sufficiently diverse. We built multiple alignments with different levels of permissiveness in including distantly related sequences, and calculated the top-scoring L/2 (L = sequence length, 56) evolutionary couplings to compare against the epistasis-derived pairings. We were only able to identify 9 high-confidence evolutionary couplings (Methods), though multiple lower-ranked couplings were true contacts in the wildtype structure (Supp. Figure 3). We see that the top-ranked positive epistatic pair is represented in this group of high-scoring evolutionary couplings (Methods, Supp. Figure 4). However, this group of evolutionary couplings does not share a strong commonality with the most positive epistatic contact predictions (Supp. Figure 4). These findings indicate that although high-ranking evolutionary couplings do have some overlap with high-ranking epistatic-based contact predictions, they provide non-identical and complementary sources of information. While the precise mechanistic details require further investigation, it is clear that the selection processes in natural evolution in many species and across many generations give rise to different details of inter-residue constraints than do screening or selection assays in the laboratory.

## Discussion

### First folding from epistatic pairs

This work was inspired by the comprehensive biophysical description of pairwise epistasis by Olson et al. (*31*) and makes use of their rich dataset screening an exhaustive library of residue pair mutants in an ingenious binding assay. Consistent with their observation that strongly epistatic pairs can be explained by contact interactions in the protein structure, we demonstrate that top-ranked epistatic pairs can be used to predict the correct protein fold to remarkable accuracy.

### Few epistatic pairs suffice

The number of epistatic pairs sufficient for folding is surprisingly small (e.g., 37 contact pairs out of 1485 residue pairs tested) and less than the 20-40% randomly sampled subsets of correct contacts needed to reach fold accuracy of 4Å-2Å positional rmsd (*50*) in benchmark tests. The reason might be that the functionally and structurally most constrained pairs occupy strategic positions for the folding process and/or the stability of the protein structure so that knowledge of only a sparse set of interaction constraints leads to the correct fold.

### Design of more efficient libraries for structure determination from pair mutation scans

As only a small subset of a complete double mutant scan carries the majority of information about pairwise proximity, it may be possible to obtain fold predictions from significantly less exhaustive experiments, if one can design – and generate at reasonable cost – a library of mutants with a higher yield of strongly epistatic pairs. Such design can be guided by various sources of information, such as prediction of secondary structure and residue surface exposure, conservation patterns from natural sequences, and physio-chemical or statistical residue-residue interaction potentials. Synthesis of designed libraries, however, is currently very costly for proteins larger than about 70 residues. A practical compromise for larger proteins may be the random ligation of designed libraries for 70-residue fragments.

### Beyond 3D structure

Similar to observations by Olson et al. *(31)*, we find that the strongest epistatic residue pairs map to near one of the two main interaction sites of the GB1 protein domain with the Fab fragment of immunoglobulin IgG (PDB ID 1IGC, PMID 7966308). These epistatic interactions appear to stabilize the interaction of the C-terminus of the α-helix with two of the beta strands and nearby loops. It is plausible that reliable formation and stability of this part of the structure is particularly important for binding in the complementary pocket on the IgG, causing a small number of strong epistatic signals in the Olsen et al. binding screen. Future scans of this type may therefore not only provide 3D folds by also indicate binding site or active sites, according to the type of assay used.

### A new experimental method for protein structure determination

Overall, high throughput selection or screening experiments systematically probing epistasis of sequence changes contain a wealth of information on interactions constrained by protein function and can provide sufficient information to determine a protein fold. Structure determination by systematic mutational scans followed by epistasis analysis is a new opportunity, complementing the classic experimental methods for protein structure determination such as crystallography, NMR or cryo-EM. To make this type of epistasis method broadly applicable for structure determination raises the challenges of the definition of assays for particular proteins, generally efficient experimental methods for reading out protein fitness for each sequence, as well as coping with much larger libraries for larger proteins.

## Methods

### Calculation of epistasis

Olson et al. synthesized a majority (535,917/536,085) of all possible double mutants, and all single mutants (1,045) in the GB1 domain of protein G (*31*). The fitness of each individual mutant was defined as the ratio of sequence reads before and after selection, normalized by the ratio for the wild-type sequence. Epistasis was defined as the logged ratio between double mutant fitness and the product of the fitness of constituent single mutants.

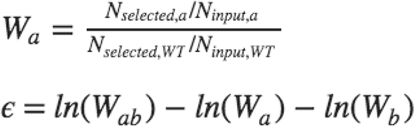

As Olson *et al*. found that non-specific absorption to Ig-G beads led to a fitness of approximately 0.01, all fitness values smaller than this were set to 0.01. Similarly, in the calculation of epistasis, when the product of single mutant fitnesses is smaller than 0.01, it is also fixed to 0.01. The input counts of double mutants vary between 1-64,627. As an additional filtering step, we excluded all mutants with fewer than 20 pre-selection read counts from analysis as lower input counts sensitize measurements to noise. This filtering step removes ~3% of the synthesized double mutants. The fitness and epistasis values of every measured double mutant can be found in Supplementary Table 1.

### Secondary structure estimation

In order to determine secondary structure from the epistasis measurements, we adapted a method from *Toth-Petroczy et al*. that predicts α and β propensities from local evolutionary couplings *(39)*. In β strands, every other position is proximal, whereas in α helices every third and fourth position is proximal. For each position i, we therefore calculated a β strand score defined by the maximum epistasis measured between position i and i+2, i-2 minus that between i and i+1, i-1. We then calculate an α score according to the maximum epistasis measured between position i and i+3, i-3, i+4, i-4. Secondary structure was then attributed wherever the mean α (from i-4 to i+4) or mean β score (from i-2 to i+2) was greater than the average over the sequence.

### 3D folding constraints and selection

To generate 3D folds of GB1 that place epistatic positions in proximity, we used the Crystallography and NMR System (CNS) package (*51*). An extended chain model was generated for GB1 and used as the starting fold. We used varying numbers of long-range (>= 5 amino acids apart in the sequence) epistasis-predicted contacts as 2-4 Å distance constraints on the most distal heavy atom of each side chain. Distance and dihedral angle constraints for the secondary structure were placed as described in previous work *(32, 33)* using the correct secondary structure regions from the crystal structure 1pga *(45)*. The standard version of CNS’s distance geometry - simulated annealing script was used to calculate 10 models for each number of top-scoring epistatic constraints between 10 and 56, and an additional level of simulated annealing based on the default <anneal.inp> script in CNS was run on each of these intermediate models using 20 or 30 epistatic constraints. This algorithm produced 4100 total folded model candidates.

### Model ranking

From the contact maps, we were able to recover the topology of the β strands, including high-scoring contacts between strand 1 and 2, 1 and 4, and 3 and 4. We ranked each model based on how many of the top L/2 (*28*) epistatic constraints were satisfied added to how many contacts between interacting strands we observed. As a control, we ranked the crystal structure and found it scored the highest compared to all of the predicted structures. The top-scoring models from this ranking were compared with the wildtype structures to report the resulting TM and rmsd scores, also finding the best ranking model by our criterion to have the lowest rmsd.

### Evolutionary couplings of GB1 from natural sequence variation

Multiple sequence alignments of the GB1 domain (Uniprot P06654, residues 225-290) were generated using five iterations of jackhammer (*52*) against the April 2017 Uniref100 dataset (*53*). Evolutionary couplings were calculated as described previously *(32, 33)* on an alignment including 59,327 sequences (14,441 effective sequences after down-weighting sequences with > 80% sequence identity) with 56/66 residues containing fewer than 30% gaps in aligned columns. A mixture model approach was used to identify residues most likely to be in the lognormal tail of the score distribution. 9 evolutionary couplings were found to have a >90% of falling in the tail and are therefore most likely to be true contacts *(39)*.

### Comparing evolutionary couplings to epistasis experiment

Positive epistasis scores were calculated by taking the most positive value over all possible mutations at each pair of residues at positions i and j, where |i-j| >= 5 to select only long-range contacts. The chance of having the highest positive epistatic score in the group of evolutionary couplings randomly was computed as 0.022 (28/N where N = 1275 possible long-range contacts, derived using the hypergeometric distribution probability of encountering our single top-ranked pair in 28 random draws over a population of 1275 without replacement.) We compared the top L/2 evolutionary couplings to a null model wherein couplings fall randomly in the epistasis distribution. Therefore, feature distributions were generated by randomly sampling groups of 28 (i,j) pairs from this set 500,000 times, and computing either the top-ranked, mean, or median epistasis value of this subset. Although the average epistatic score for the top L/2 evolutionary couplings was greater than 96% of random groups of pairs of the same size, the median of the evolutionary couplings group was indistinguishable from the distribution of medians from random samples (Supp. Figure 4).

When classifying contacts as unique in one fold versus common in another, we require atoms in one residue to be within 5Å of any atom in another residue but also more than 8Å apart from that residue in the alternative structure.

## Acknowledgements

All authors thank the Marks and Sander labs for discussion and support, and NR also thanks the Silver lab for discussions and support.

While completing this project, we became aware of a related effort by J. Schmiedel and B. Lehner who also report the prediction of good accuracy 3D structures of the GB1 domain based on analysis of epistasis patterns in the Olson et al. mutation scan (bioRxiv 303875; doi: https://doi.org/10.1101/303875).

## Supplementary information

**Supplementary figure 1:**
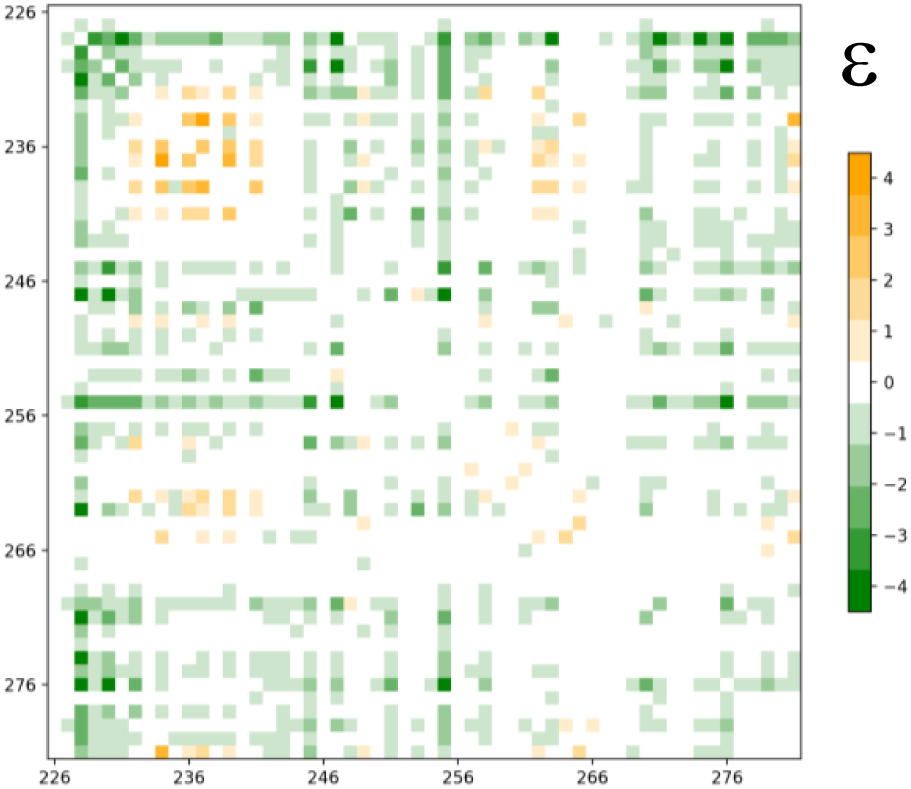
Mean epistasis for all mutations at positions i,j. Values scaled by 3 to meet the colorscale used by *Olson et al*. Figure 3b. More negative average epistasis is shown as green, more positive as orange.

**Supplementary figure 2:**
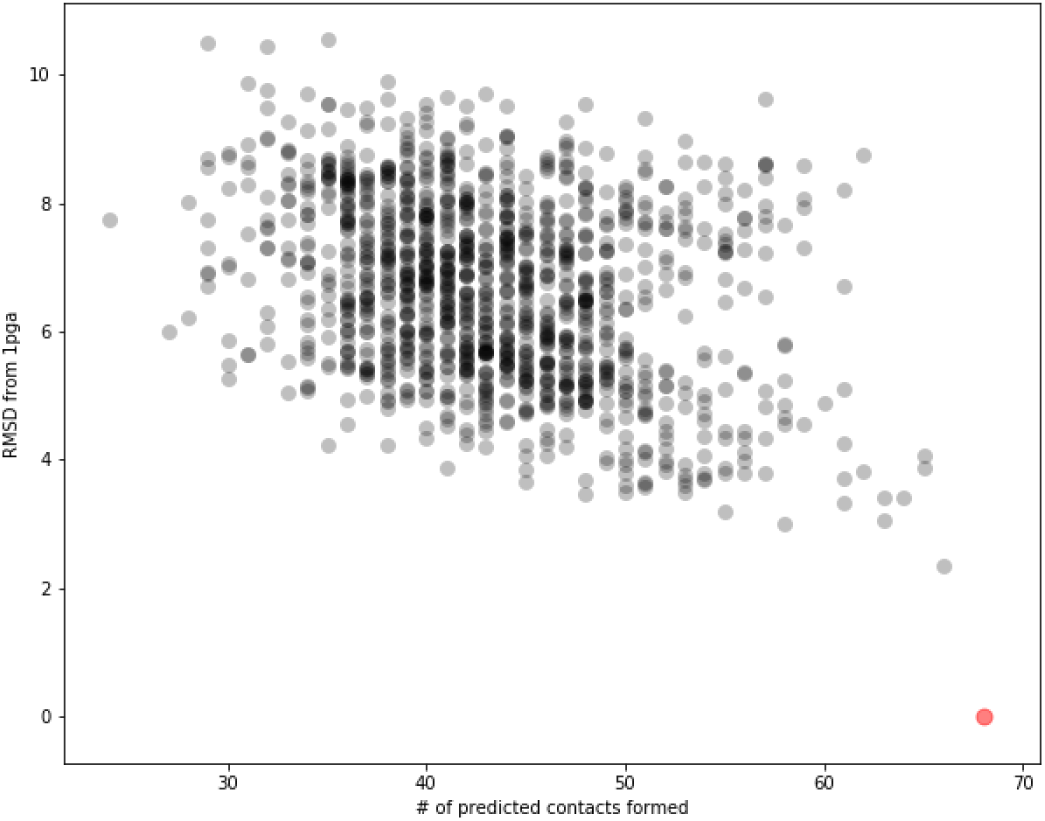
The root mean square distance of backbone carbons in the CNS models to those of a gb1 crystal (pdb id: 1pga), plotted versus ranking score. The 150 constraints are obtained from the top L/2 epistasis pairs, and all possible interactions between the beta strands implicated by the secondary structure and topology inferred by epistasis-predicted contacts (see Figure 1). 1pga is shown in red, with 0 rmsd, satisfying 68 constraints.

**Supplementary figure 3:**
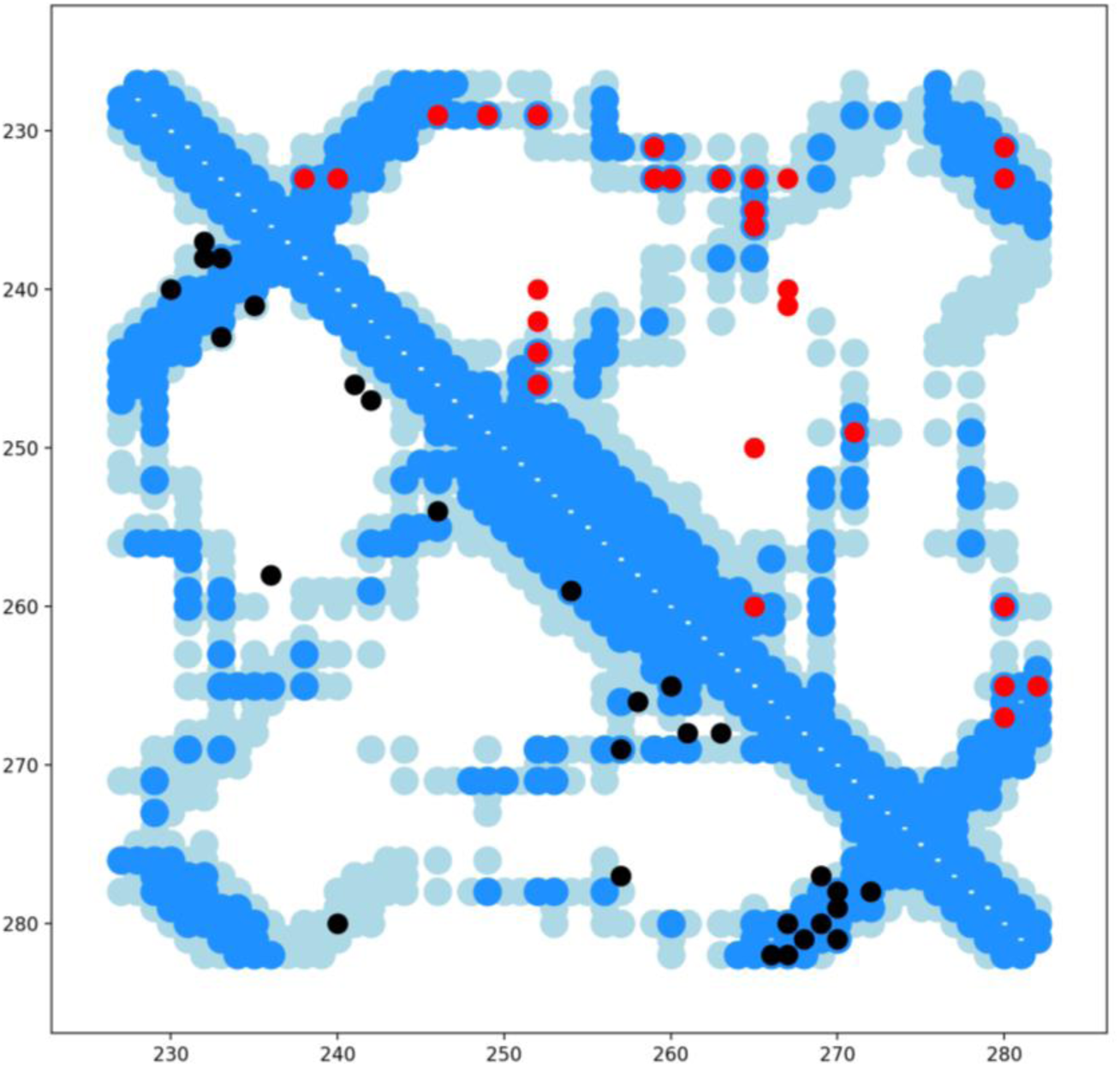
Top L/2 evolutionary couplings (black) and top L/2 positive epistasis pairs (red), plotted on top of the contacts (<5A, dark blue; <8A, light blue) of a gb1 structure (pdb id: 2gb1). To differentiate the couplings from epistatic pairs, lower and upper triangles are plotted respectively.

**Supplementary figure 4:**
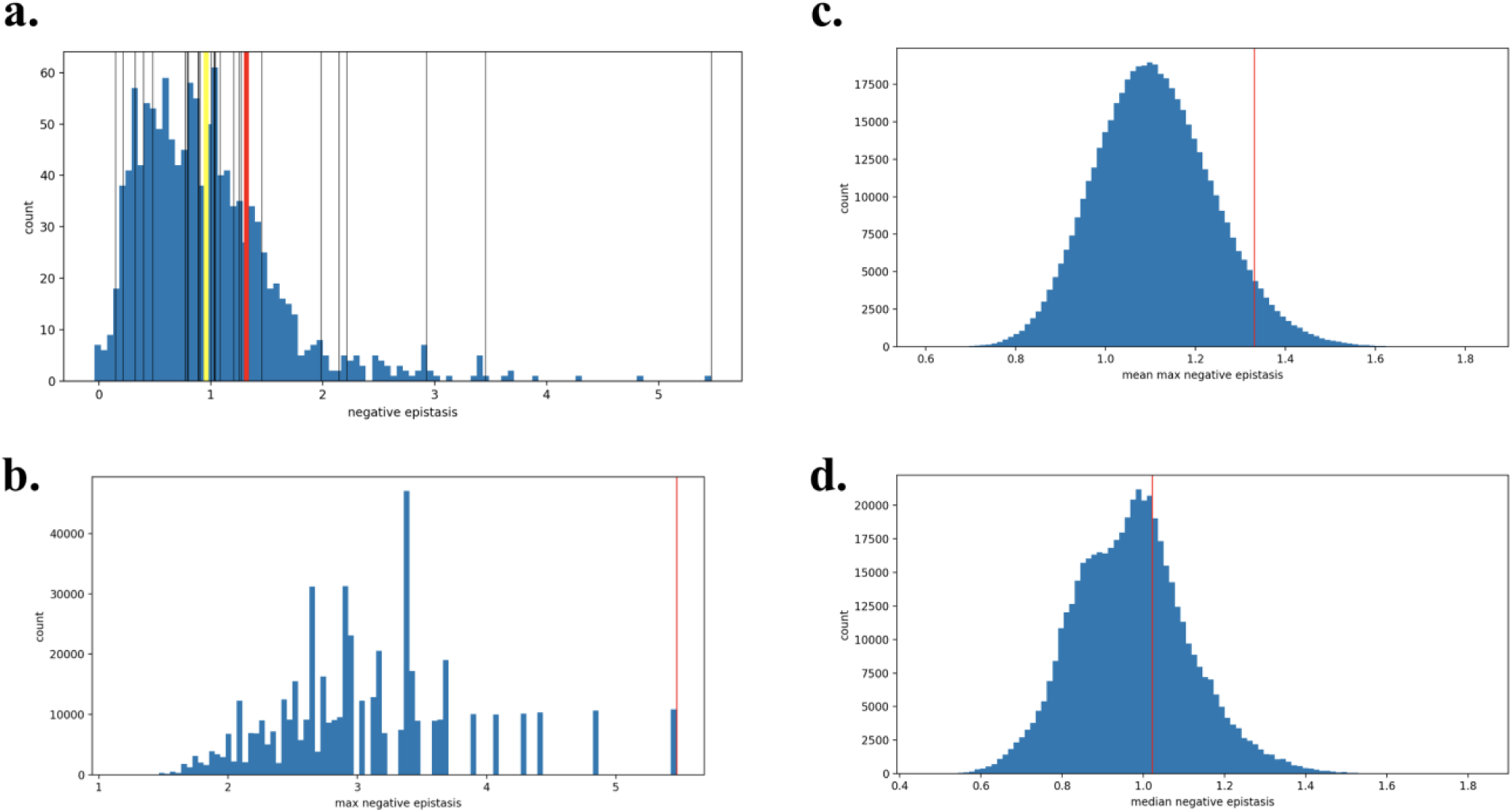
The top L/2 evolutionary couplings compared against the largest values of positive epistasis at all pairs of positions i,j. **a**. The measured max positive epistasis at each of the top L/2 couplings (black) and the mean of those values (red) versus the overall distribution of max positive epistasis across all i,j-s (blue) with mean (yellow). **b**. The distribution of max positive epistasis for 500,000 samplings of 28 (L/2) random i,j pairs (blue). The max positive epistasis value among the evolutionary couplings (red). **c**. The distribution of mean positive epistasis values (blue) and mean of the evolutionary couplings (red). **d**. The distribution of median positive epistasis values (blue) and median of the evolutionary couplings (red).

**Supplementary table 1:**

Calculated fitness and epistasis values for all double mutants-raw input and output counts are included from *Olson et al*.

**Supplementary table 2:**

True positives of the top positive, negative and absolute epistasis pairs versus all <5A crystal contacts from wild-type and mutant gb1.

**Supplementary table 3:**

Alpha and beta secondary structure scores computed from epistasis values, as described in Methods.

## References

1. J. Chen, W. E. Stites, Energetics of side chain packing in staphylococcal nuclease assessed by systematic double mutant cycles. Biochemistry 40, 14004–14011 (2001).

2. E. J. Ackermann, E. T. Ang, J. R. Kanter, I. Tsigelny, P. Taylor, Identification of pairwise interactions in the alpha-neurotoxin-nicotinic acetylcholine receptor complex through double mutant cycles. J Biol Chem 273, 10958–10964 (1998).

3. A. Horovitz, Double-mutant cycles: a powerful tool for analyzing protein structure and function. Fold Des 1, R121–126 (1996).

4. P. A. Romero, T. M. Tran, A. R. Abate, Dissecting enzyme function with microfluidic-based deep mutational scanning. Proc Natl Acad Sci U S A 112, 7159–7164 (2015).

5. B. P. Roscoe, D. N. Bolon, Systematic exploration of ubiquitin sequence, E1 activation efficiency, and experimental fitness in yeast. Journal of molecular biology 426, 2854–2870 (2014).

6. B. P. Roscoe, K. M. Thayer, K. B. Zeldovich, D. Fushman, D. N. Bolon, Analyses of the effects of all ubiquitin point mutants on yeast growth rate. Journal of molecular biology 425, 1363–1377 (2013).

7. D. Melamed, D. L. Young, C. E. Gamble, C. R. Miller, S. Fields, Deep mutational scanning of an RRM domain of the Saccharomyces cerevisiae poly(A)-binding protein. Rna 19, 1537–1551 (2013).

8. M. A. Stiffler, D. R. Hekstra, R. Ranganathan, Evolvability as a Function of Purifying Selection in TEM-1 beta-Lactamase. Cell 160, 882–892 (2015).

9. R. N. McLaughlin, Jr., F. J. Poelwijk, A. Raman, W. S. Gosal, R. Ranganathan, The spatial architecture of protein function and adaptation. Nature 491, 138–142 (2012).

10. J. O. Kitzman, L. M. Starita, R. S. Lo, S. Fields, J. Shendure, Massively parallel single-amino-acid mutagenesis. Nature methods 12, 203–206, 204 p following 206 (2015).

11. A. Melnikov, P. Rogov, L. Wang, A. Gnirke, T. S. Mikkelsen, Comprehensive mutational scanning of a kinase in vivo reveals substrate-dependent fitness landscapes. Nucleic acids research 42, e112 (2014).

12. C. L. Araya et al., A fundamental protein property, thermodynamic stability, revealed solely from large-scale measurements of protein function. Proc Natl Acad Sci U S A 109, 16858–16863 (2012).

13. E. Firnberg, J. W. Labonte, J. J. Gray, M. Ostermeier, A comprehensive, high-resolution map of a gene’s fitness landscape. Mol Biol Evol 31, 1581–1592 (2014).

14. L. M. Starita et al., Massively Parallel Functional Analysis of BRCA1 RING Domain Variants. Genetics, (2015).

15. L. Rockah-Shmuel, A. Toth-Petroczy, D. S. Tawfik, Systematic Mapping of Protein Mutational Space by Prolonged Drift Reveals the Deleterious Effects of Seemingly Neutral Mutations. PLoS Comput Biol 11, e1004421 (2015).

16. H. Jacquier et al., Capturing the mutational landscape of the beta-lactamase TEM-1. Proc Natl Acad Sci U S A 110, 13067–13072 (2013).

17. H. Qi et al., A quantitative high-resolution genetic profile rapidly identifies sequence determinants of hepatitis C viral fitness and drug sensitivity. PLoS Pathog 10, e1004064 (2014).

18. N. C. Wu et al., Functional Constraint Profiling of a Viral Protein Reveals Discordance of Evolutionary Conservation and Functionality. PLoS Genet 11, e1005310 (2015).

19. P. Mishra, J. M. Flynn, T. N. Starr, D. N. Bolon, Systematic Mutant Analyses Elucidate General and Client-Specific Aspects of Hsp90 Function. Cell Rep 15, 588–598 (2016).

20. M. B. Doud, J. D. Bloom, Accurate measurement of the effects of all amino-acid mutations to influenza hemagglutinin. *bioRxiv*, (2016).

21. Z. Deng et al., Deep sequencing of systematic combinatorial libraries reveals beta-lactamase sequence constraints at high resolution. Journal of molecular biology 424, 150–167 (2012).

22. L. M. Starita et al., Activity-enhancing mutations in an E3 ubiquitin ligase identified by high-throughput mutagenesis. Proc Natl Acad Sci U S A 110, E1263–1272 (2013).

23. C. D. Aakre et al., Evolving new protein-protein interaction specificity through promiscuous intermediates. Cell 163, 594–606 (2015).

24. P. Julien, B. Minana, P. Baeza-Centurion, J. Valcarcel, B. Lehner, The complete local genotype-phenotype landscape for the alternative splicing of a human exon. Nat Commun 7,11558 (2016).

25. C. Li, W. Qian, C. J. Maclean, J. Zhang, The fitness landscape of a tRNA gene. Science, (2016).

26. D. Mavor et al., Determination of ubiquitin fitness landscapes under different chemical stresses in a classroom setting. Elife 5, (2016).

27. D. M. Fowler, S. Fields, Deep mutational scanning: a new style of protein science. Nature methods 11, 801–807 (2014).

28. M. Gasperini, L. Starita, J. Shendure, The power of multiplexed functional analysis of genetic variants. Nat Protoc 11, 1782–1787 (2016).

29. L. M. Starita et al., Variant Interpretation: Functional Assays to the Rescue. Am J Hum Genet 101, 315–325 (2017).

30. K. S. Sarkisyan et al., Local fitness landscape of the green fluorescent protein. Nature 533, 397–401 (2016).

31. C. A. Olson, N. C. Wu, R. Sun, A comprehensive biophysical description of pairwise epistasis throughout an entire protein domain. Current biology: CB 24, 2643–2651 (2014).

32. T. A. Hopf et al., Three-Dimensional Structures of Membrane Proteins from Genomic Sequencing. Cell 149, 1607–1621 (2012).

33. D. S. Marks et al., Protein 3D Structure Computed from Evolutionary Sequence Variation. PLOS ONE 6, e28766 (2011).

34. F. Morcos et al., Direct-coupling analysis of residue coevolution captures native contacts across many protein families. Proc Natl Acad Sci U S A 108, E1293–1301 (2011).

35. T. Kosciolek, D. T. Jones, De novo structure prediction of globular proteins aided by sequence variation-derived contacts. PLoS One 9, e92197 (2014).

36. S. Ovchinnikov et al., Large-scale determination of previously unsolved protein structures using evolutionary information. Elife 4, e09248 (2015).

37. T. A. Hopf et al., Sequence co-evolution gives 3D contacts and structures of protein complexes. Elife 3, (2014).

38. C. Weinreb et al., 3D RNA and Functional Interactions from Evolutionary Couplings. Cell 165, 963–975 (2016).

39. A. Toth-Petroczy et al., Structured States of Disordered Proteins from Genomic Sequences. Cell 167, 158–170.e112 (2016).

40. P. A. Alexander, Y. He, Y. Chen, J. Orban, P. N. Bryan, A minimal sequence code for switching protein structure and function. Proc Natl Acad Sci U S A 106, 21149–21154 (2009).

41. Y. He, Y. Chen, P. Alexander, P. N. Bryan, J. Orban, NMR structures of two designed proteins with high sequence identity but different fold and function. Proc Natl Acad Sci U S A 105, 14412–14417 (2008).

42. Y. He, Y. Chen, P. A. Alexander, P. N. Bryan, J. Orban, Mutational tipping points for switching protein folds and functions. Structure 20, 283–291 (2012).

43. C. Falkenberg, L. Bjorck, B. Akerstrom, Localization of the binding site for streptococcal protein G on human serum albumin. Identification of a 5.5-kilodalton protein G binding albumin fragment. Biochemistry 31, 1451–1457 (1992).

44. A. M. Gronenborn et al., A novel, highly stable fold of the immunoglobulin binding domain of streptococcal protein G. Science 253, 657–661 (1991).

45. T. Gallagher, P. Alexander, P. Bryan, G. L. Gilliland, Two crystal structures of the B1 immunoglobulin-binding domain of streptococcal protein G and comparison with NMR. Biochemistry 33, 4721–4729 (1994).

46. G. Yachdav et al., PredictProtein--an open resource for online prediction of protein structural and functional features. Nucleic acids research 42, W337–343 (2014).

47. D. W. Buchan, F. Minneci, T. C. Nugent, K. Bryson, D. T. Jones, Scalable web services for the PSIPRED Protein Analysis Workbench. Nucleic acids research 41, W349–357 (2013).

48. A. T. Brunger, Version 1.2 of the Crystallography and NMR system. Nat Protoc 2, 2728–2733 (2007).

49. Y. Zhang, J. Skolnick, TM-align: a protein structure alignment algorithm based on the TM-score. Nucleic Acids Res 33, 2302–2309 (2005).

50. R. Sathyapriya, J. M. Duarte, H. Stehr, I. Filippis, M. Lappe, Defining an essence of structure determining residue contacts in proteins. PLOS Comput Biol 5, e1000584 (2009).

51. A. T. Brünger et al., Crystallography & NMR system: A new software suite for macromolecular structure determination. Acta Crystallogr. D Biol. Crystallogr. 54, 905–921 (1998).

52. L. S. Johnson, S. R. Eddy, E. Portugaly, Hidden Markov model speed heuristic and iterative HMM search procedure. BMC Bioinformatics 11, 431 (2010).

53. B. E. Suzek, Y. Wang, H. Huang, P. B. McGarvey, C. H. Wu, UniRef clusters: a comprehensive and scalable alternative for improving sequence similarity searches. Bioinformatics 31, 926–932 (2015).

54. A. E. Sauer-Eriksson, G. J. Kleywegt, M. Uhlen, T. A. Jones, Crystal structure of the C2 fragment of streptococcal protein G in complex with the Fc domain of human IgG. Structure 3, 265–278 (1995).

